# Understanding the antiviral effects of RNAi-based therapy on chronic hepatitis B infection

**DOI:** 10.1101/2020.07.21.215012

**Authors:** Sarah Kadelka, Harel Dahari, Stanca M Ciupe

## Abstract

Reaching hepatitis B surface antigen (HBsAg) loss (called functional cure) with approved treatment with pegylated interferon-*α* (IFN) and/or nucleos(t)ide analogues (NAs) in chronic hepatitis B virus (HBV) infected patients is suboptimal. The RNA interference (RNAi) drug ARC-520 was shown to be effective in reducing serum HBV DNA, HBsAg and hepatitis B e antigen (HBeAg) in chimpanzees and small animals. A recent clinical study (Heparc-2001) showed reduction of serum HBV DNA, HBeAg and HBsAg in HBeAg-positive patients treated with a single dose of ARC-520 and daily NA (entecavir). To provide insights into HBV dynamics under ARC-520 treatment and its efficacy in blocking HBV DNA, HBsAg, and HBeAg production we developed a a multi-compartmental pharmacokinetic-pharamacodynamic model and calibrated it with measured HBV data. We showed that the time-dependent ARC-520 efficacies in blocking HBsAg and HBeAg are more than 96% effective around day 1, and slowly wane to 50% in 1-4 months. The combined ARC-520 and entecavir effect on HBV DNA is constant over time, with efficacy of more than 99.8%. HBV DNA loss is entecavir mediated and the strong but transient HBsAg and HBeAg decays are solely ARC-520 mediated. We added complexity to the model in order to reproduce current long-term therapy outcomes with NAs by considering the tradeoff between hepatocyte loss and hepatocyte division, and used it to make *in-silico* long-term predictions for virus, HBsAg and HBeAg titer dynamics. These results may help assess ongoing RNAi drug development for hepatitis B virus infection.

**Author summary:** With about 300 million persons infected worldwide and 800,000 deaths annually, chronic infection with hepatitis B virus (HBV) is a major public health burden with high endemic areas around the world. Current treatment options focus on removing circulating HBV DNA but are suboptimal in removing hepatitis B s- and e-antigens. ARC-520, a RNA interference drug, had induced substantial hepatitis B s- and e-antigen reductions in animals and patients receiving therapy. We study the effect of ARC-520 on hepatitis B s- and e-antigen decline by developing mathematical models for the dynamics of intracellular and serum viral replication, and compare it to patient HBV DNA, hepatitis B s- and e-antigen data from a clinical trial with one ARC-520 injection and daily nucleoside analogue therapy. We examine biological parameters describing the different phases of HBV DNA, s-antigen and e-antigen decline and rebound after treatment initiation, and estimate treatment effectiveness. Such approach can inform the RNA interference drug therapy.

## Introduction

Treatment options for chronic hepatitis B (HBV) infections are limited to two main drug groups: pegylated interferon-*α* (IFN) and nucleos(t)ide analogues (NAs) [1–3]. Treatment with IFN induces antiviral activity, immunomodulatory effects, and robust off-treatment responses. These responses, however, vary among patients and induce *functional cure*, defined as hepatitis B surface antigen (HBsAg) loss, in only 10 − 20% Caucasian patients and less than 5% Asian patients. Moreover, IFN treatment is poorly tolerated [4–6]. By contrast, treatment with NAs is well tolerated and can be life-long but has limited effect in reducing serum HBsAg and hepatitis B e-antigen (HBeAg) production and, in limiting hepatitis B covalently closed circular DNA (cccDNA) persistence and HBV DNA integration [1, 7, 8], all of which play important roles in chronic infections. HBeAg is thought to induce T cell tolerance to both e- and core antigens and to be an important reason for viral persistence [9]. HBsAgs, besides being used for virion envelopes, form empty non-infectious subviral particles (*i.e.* without viral genome) whose numbers are at least 1,000-folds higher than those of virions [10], and may serve as decoy for antibody responses [11]. Moreover, they are also assumed to be involved in T cell exhaustion [12, 13]. Functional cure has been proposed as a desirable outcome of treatment. None of the currently licensed therapies can produce this result for a large fraction of chronically infected patients. There is therefore a need for new therapies that target HBsAg production and/or its clearance from circulation [14, 15].

RNA interference (RNAi) technology has the ability of silencing specific genes and can, therefore, be used for treatment against a large array of infectious agents (see [16] for a review on RNAi-based therapies). For hepatitis B infection, small interfering RNAs were designed to hybridize with HBV mRNA inside an infected hepatocyte and, as a result, induce its degradation [17, 18]. ARC-520, the first such small interfering RNA to be tested in clinical trials, was designed with the aim of knocking down the expression of all HBV mRNA, including HBsAg proteins. Experiments in mice and chimpanzees, and a phase II clinical study in patients (Heparc-2001) showed potential for ARC-520 induced HBeAg, HBsAg and HBV DNA titers reduction [17, 19]. The Heparc-2001 study showed differential HBsAg reduction among patients based on their HBeAg status and prior exposure to traditional therapy such as NAs [19]. While ARC-520 has been terminated due to delivery-associated toxicity [19], overall results indicate that RNAi-based therapy has the potential of reducing HBsAg and inducing functional cure. Therefore, a next generation of RNAi drugs with improved delivery methods may serve as means for protein removal and HBV functional cure. Two such RNAi therapies are currently undergoing clinical trials with promising results [16].

To better understand the effect of RNAi therapies, additional information regarding the host-virus-drug dynamics and therapy outcomes are needed. In this study, we developed mathematical models that best reproduce observed HBV DNA, HBsAg and HBeAg kinetics following a single dose of ARC-520 in five HBeAg-positive patients from the Heparc-2001 study. Mathematical models of hepatitis B infection have been used to study the dynamics of acute, chronic, and occult HBV infections [20–24], anti-HBV therapy [14, 25–30], cell-to-cell transmission [31], intracellular interactions [31–33], cellular immune responses [21, 25, 34–36], antibody-mediated immune responses [11, 33, 37], HBeAg [33, 38, 39], and HBeAb [33] dynamics. We build on previous modeling work, consider the interaction between HBV DNA, HBsAg and HBeAg titers in the presence of a single dose RNAi-based therapy, and use the model to run *in silico* experiments to predict individual contributions of different drug effects on the dynamics for HBsAg titers.

## Methods

### Patient data

We use published data from five HBeAg-positive, treatment-naive chronic hepatitis B patients (cohort 7 in [19]), which are the ones that best responded to ARC-520 therapy. Moreover, they are the only studied cohort in which HBV DNA integration is not reported as a source of HBsAg production (as opposed to HBeAg-negative and NA-experienced HBeAg-positive patients with low cccDNA), and, thus, it allowed us to exclude integration when developing the mathematical model. Data consists of serum HBV DNA titers (in IU/ml), HBsAg, and HBeAg concentration (in IU/ml) measured at *t*_*i*_= {−8, 0, 2, 7, 14, 21, 28, 42, 56, 84} days, where *i* = {−1, *…*, 8} and *t*_0_ = 0 is the day when both daily NA entecavir (ETV) and a single intravenous ARC-520 injection (inoculum of 4 mg/kg) are administrated.

### Pharmacokinetics-pharamcodynamics model

We are interested in determining the mechanisms underlying the observed HBV DNA, HBsAg and HBeAg kinetics under combined ETV and ARC-520 therapy. We develop a mathematical model that considers the interactions between infected hepatocytes, *I* (in cells per ml); total intracellular HBV DNA, *D* (in copies per ml); serum HBV DNA, *V* (in IU per ml); serum HBsAg, *S* (in IU per ml); and serum HBeAg, *E* (in IU per ml). We assume that infected cells decay at per capita rate *δ*, and we exclude cell proliferation (we will relax this assumption later on). We assume intracellular HBV DNA is synthesized at rate *α* and is lost at constant per capita rate *c*_*D*_. The replication rate *α* summarizes various steps that are not modeled explicitly, such as the transcription of pregenomic RNA (pgRNA) from cccDNA, and the generation of single stranded DNA by reverse transcription. Intracellular HBV DNA is assembled and released into blood as free virions at rate *p* which are cleared at rate *c* [40]. To account for the different units of intracellular and serum virus, we use the conversion factor *ξ* = 1*/*5.3 IU/copies [41]. Lastly, we assume HBsAg and HBeAg are transcribed from cccDNA inside infected hepatocytes and then released into blood at rates *p*_*S*_ and *p*_*E*_, respectively, and are cleared at per capita rates *d*_*S*_ and *d*_*E*_, respectively. We have not included HBV DNA integration, which is only a substantial source of HBsAg in HBeAg negative patients and NUC-experienced HBeAg positive patients with low cccDNA [19]. The model is given by the following model:

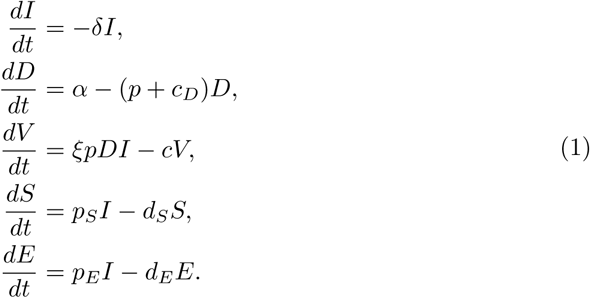

Patients were administered daily nucleoside analogous treatment with entecavir starting at day *t*_0_ = 0. ETV is known to block reverse transcription of HBV DNA, and therefore inhibit HBV DNA synthesis. We model this (see model (5)) as a constant reduction of the HBV DNA synthesis rate *α* to (1 − *ϵ*)*α*, where 0 ≤ *ϵ* ≤ 1 is the ETV efficacy. Experimental studies in humanized mice have shown that serum HBV DNA declines in biphasic manner while HBV-infected cell are not lost in the first months following NA treatment initiation [42]. To account for the biphasic HBV DNA decay in the absence of infected cell killing, we assume that ETV has additional time-dependent inhibitory effects on intracellular HBV DNA synthesis and model it by decreasing *α* further to 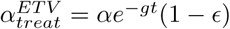, where *g* ≥ 0 is a constant and *t* is the time in days post ETV initiation. Moreover, a single ARC-520 dose was administrated at time *t*_0_ = 0. Unlike ETV, which was given daily, we model the build-up and clearance of ARC-520 pharmacokinetics over time by considering a two-compartment pharmacokinetic model consisting of drug quantity in the plasma and liver, *C*_*p*_ and *C*_*e*_, respectively [43]. The inoculum *C*_*p*_(0) = *C*_0_ decays exponentially at rate 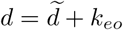, where 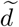 is the plasma drug degradation rate and *k*_*eo*_ is the absorption into the liver rate. The drug in the liver decays at rate *k*_*eo*_, identical with the absorption rate [44]. Following these assumptions, the pharmacokinetic model has the form:

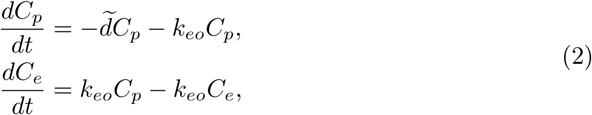

with initial conditions *C*_*p*_(0) = *C*_0_ and *C*_*e*_(0) = 0. This is a linear model which can be solved to give solutions:

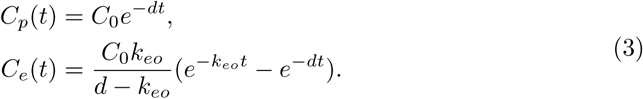

Lastly, we assume the relationship between the drug quantity in the liver *C*_*e*_(t) and drug efficacy *η*_*i*_(*t*) to be given by:

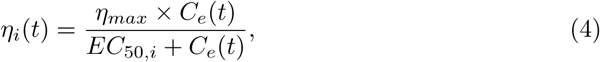

where *η*_*max*_ = 1 is the maximum drug efficacy, *EC*_50,*i*_ are drug quantities that yield half-maximal effects, and *i* = {1, 2, 3} are the infectious events that are affected by ARC-520 therapy, *i.e.*, the transcription of HBV DNA, the transcription of HBsAg, and the transcription of HBeAg, respectively. The effects of ARC-520 on intracellular HBV DNA, HBsAg and HBeAg are modeled as the reduction of intracellular HBV DNA synthesis *α* to 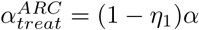, HBsAg production from *p*_*S*_ to *p*_*S,treat*_ = (1 − *η*_2_)*p*_*S*_, and of HBeAg production from *p*_*E*_ to *p*_*E,treat*_ = (1 − *η*_3_)*p*_*E*_, respectively. Considered together, models (1) and (4) give the following pharmacokinetics-pharamcodynamics (PK/PD) model:

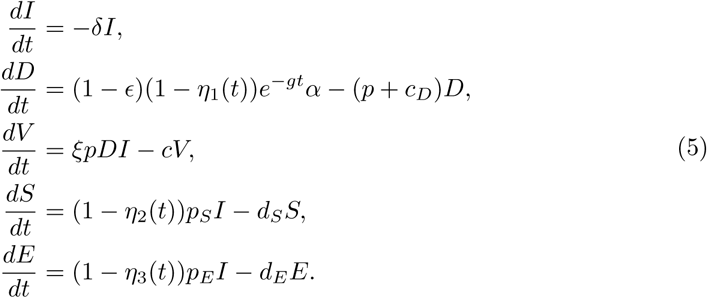

### Data fitting

We used published kinetic HBV DNA, HBsAg, HBeAg data in serum measured from five HBeAg-positive, treatment-naive chronic hepatitis B patients as described in the ‘Patient data’ section.

#### Parameter values

We assume that, prior to therapy initiation, model (5) describes a persistent chronic infection and is at the quasi-equilibrium, given by the initial values *I*(0) = *I*_0_, *D*(0) = *D*_0_, *V* (0) = *V*_0_, *S*(0) = *S*_0_ and *E*(0) = *E*_0_. Initial values for HBV DNA, *V* (0) = *V*_0_; HBsAg, *S*(0) = *S*_0_; and HBeAg, *E*(0) = *E*_0_, are set to the patient data prior to the start of therapy, *t*_−1_ = − 8, (day eight prior to the ARC-520 injection). The percentage of HBV-infected hepatocytes is reported to vary between 18 ± 12% in chronic HBsAg carriers [45, 46] and 99% in acute infections [21, 47]. Without loss of generality, we arbitrary assume that 50% of hepatocytes are infected at the beginning of treatment. Liver contains approximately 2 × 10^11^ hepatocytes, which, when distributed throughout 15 liters of extracellular fluid, gives a total hepatocyte concentration *T*_*max*_ = 1.4 × 10^7^ cells/ml [48]. We set the initial infected hepatocyte population to *I*_0_ = 0.5*T*_*max*_. Lastly, the pre-treatment level of intracellular HBV DNA in HBeAg positive patients is set to *D*_0_ = 225*/*(*I*_0_*/T*_*max*_) = 450 copies/ infected cell, as in [49].

Since we assume that model (5) is in chronic equilibrium (for the additional assumption *δ* = 0) before the therapy initiation, parameters *α, p, p*_*S*_, *p*_*E*_ are fixed according to the following formulas:

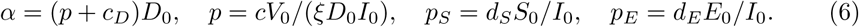

We start by ignoring the dynamics of infected cells, such as infection of susceptible cells and/or infected cell proliferation (we will relax this assumption in later sections), and assume that infected cells decay due to natural death and immune mediated killing at per capita rate *δ* = 4 × 10^−3^ per day, corresponding to a life-span of 250 days (we will later investigate the effect of increasing the killing rate, to include increased immune mediated killing or RNAi induced toxicity and death). The estimated half-life of intracellular HBV DNA is 24 hours [40, 50], which corresponds to the intracellular HBV DNA decay rate *c*_*D*_ = 0.69 per day. ARC-520’s half-life has been reported to range between 3 and 5 hours [51], corresponding to decay rates 3.3 *< d <* 5.5 per day; we fix *d* = 4 per day. Lastly, we set the initial ARC-520 quantity to the trial dose of *C*_0_ = 4 mg/kg.

The unknown parameters are **parm** = {*g, c, d*_*S*_, *d*_*E*_, *ϵ*_*T*_, *EC*_2_, *EC*_3_, *k*_*eo*_}. Here, (1 − *ϵ*_*T*_) = (1 − *ϵ*)(1 − *η*_1_(*t*)) accounts for the total drug effect on HBV DNA production. Since preliminary simulations (not shown) indicate that *η*_1_(*t*) is time independent, we cannot separate the ETV effects 1 − *ϵ* from the ARC-520 effects 1 − *η*_1_(*t*). We lump them together, and assume a total drug effect, which ranges between 0.9 *< ϵ*_*T*_ *<* 1. The other parameter ranges are found as follows. The time-dependent inhibitory effects of treatment on intracellular HBV DNA production, *g*, was estimated from HBV infected humanized mice treated with NA to range between 0.059 and 0.42 per day [40]. We expand this range by searching over the parameter space 0 *< g <* 1. There is a wide range of estimates for the free virus clearance rate in serum: as low as 0.69 per day [20, 28, 52]; and as high as 21.7 per day [53]; we search the entire 0 *< c <* 100 parameter space. The decay rate of HBsAg is bounded between 0 *< d*_*S*_ *<* 200 per day, containing previous estimates ranging between 0.057 to 0.58 per day [54, 55]. In previous modeling work [39, 56] HBeAg decay rate *d*_*E*_ was set to 0.3 per day. We allow for a larger range 0 *< d*_*E*_ *<* 200 per day, corresponding to half-lives greater than 5 minutes. We assume that the drug absorption rate *k*_*eo*_ ranges between 0 *< k*_*eo*_ *<* 1 per day. Since ARC-520 was reported to have long lasting effects [51], we assume a large range for the half-maximal quantity *EC*_*i*_; between 10^−7^ *< EC*_*i*_ *<* 1 mg/kg. These ranges are summarized in table 1.

**Table 1.** Variables and parameters in model (5). Parameters indicated by a * are fitted within the given range.

| Variables | Description | Units |  | Initial values |
| --- | --- | --- | --- | --- |
| $I$ | infected hepatocytes | cells/ml | | $0.7 \times 10^6$ |
| $D$ | intracellular HBV DNA | copies/cell | | 450 [49] |
| $V$ | free virions | IU/ml | | data at time $t_{-1} = -8$ |
| $S$ | serum HBsAg | IU/ml | | data at time $t_{-1} = -8$ |
| $E$ | serum HBeAg | IU/ml | | data at time $t_{-1} = -8$ |
| Parameters | Descriptions | Units | Default values / range | Reference |
| $\delta$ | infected cells decay rate | 1/day | $4 \times 10^{-3}$ | |
| $g^*$ | inhibitory effects on intracellular HBV production during treatment | 1/day | [0, 1] | |
| $\alpha$ | intracellular HBV DNA synthesis rate | copies/(cell×day) | $(p + c_D)D_0$ | |
| $c_D$ | intracellular HBV DNA decay rate | 1/day | 0.69 | [50] |
| $\xi$ | conversion factor | IU/copies | 1/5.3 | [41] |
| $p$ | intracellular HBV DNA release rate | 1/day | $cV_0/(\xi D_0 I_0)$ | |
| $c^*$ | free virion clearance rate | 1/day | [0, 100] | |
| $p_S$ | HBsAg production rate | IU/(cell×day) | $d_S S_0/I_0$ | |
| $p_E$ | HBeAg production rate | IU/(cell×day) | $d_E E_0/I_0$ | |
| $d_S^*$ | HBsAg decay rate | 1/day | [0, 200] | |
| $d_E^*$ | HBeAg decay rate | 1/day | [0, 200] | |
| $\epsilon_T^*$ | combined ETV and ARC-520 efficacy | unitless | [0.9,1] | |
| $C_0$ | initial plasma drug quantity | mg/kg | 4 | [19] |
| $d$ | ARC-520 decay rate | 1/day | 4 | [51] |
| $EC_2^*$ | ARC-520 quantity where $\eta_2$ is half maximal | mg/kg | $[10^{-7}, 1]$ | |
| $EC_3^*$ | ARC-520 quantity where $\eta_3$ is half maximal | mg/kg | $[10^{-7}, 1]$ | |
| $k_{eo}^*$ | drug absorption rate | 1/day | [0,1] | |

#### Optimization algorithm

We estimate the unknown parameters **parm** given in table 1 by minimizing the least squares functional:

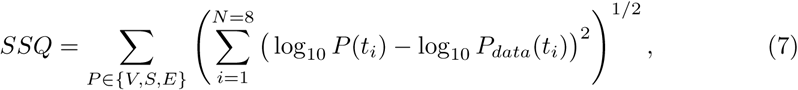

for each patient. Functional *SSQ* describes the distance between HBV DNA, HBsAg, and HBeAg titers *V*_*data*_(*t*_*i*_), *S*_*data*_(*t*_*i*_), *ϵ*_*data*_(*t*_*i*_) at times *t*_*i*_ (*i* = {1, *…*, 8}) and populations *V* (*t*_*i*_), *S*(*t*_*i*_) and *E*(*t*_*i*_) as given by model (5) at times *t*_*i*_ (*i* = {1, *…*, 8}). As described previously (see eq (6)), the before treatment titers at *t*_−1_ = −8 days are used to determine parameters *α, p, p*_*S*_, *p*_*E*_ such that the model’s equilibrium matches the titers exactly. Since we assume that the model stays in equilibrium until treatment initiation, we ignore the titers at time *t*_0_ = 0 days. Lastly, it should be noted that we assign the same weight to errors in HBV DNA, HBsAg, and HBeAg. Within the parameter space defined in table 1, we determine optimal parameter fits for each patient by following four steps (code available upon publication):

1. We create 100 parameter sets using the Latin hypercube samples (LHS) *Matlab* routine *lhsdesign*, with random number generator seed two and uniform probability density distribution on each parameter interval. Since the parameter space spans several orders of magnitude in *EC*_2_ and *EC*_3_ directions, we replace them with 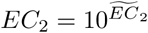 and 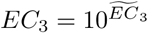. Thus, instead of sampling *EC*_2_ and *EC*_3_ in [10^−7^, 1], we sample 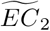 and 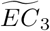 in [−7, 0]. Our preliminary work showed that *ϵ*_*T*_ ≈ 1 often yields the best results.Therefore, we replace 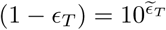 and sample 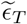 in the parameter space [−8, −1].
2. HBV DNA dynamics do not influence HBsAg and HBeAg dynamics. Therefore, we minimize 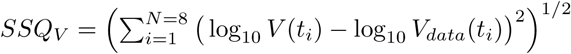 and 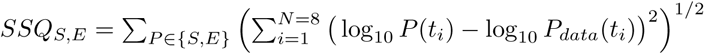 separately over their corresponding parameter sets **parm**_*V*_ = {*g, c, ϵ*_*T*_ } and **parm**_*SE*_ = {*d*_*S*_, *d*_*E*_, *EC*_2_, *EC*_3_, *k*_*eo*_}, respectively. We split the LHS into LHS_*V*_ and LHS_*S,E*_ containing the respective initial parameter guesses and, using *Matlab*’s *fmincon* routine to minimize *SSQ*_*V*_ and *SSQ*_*S,E*_ within the parameter space in table 1, obtain 100 *optimal* **parm**_*V*_ and **parm**_*S,E*_ parameter sets.
3. Of the 2×100 *optimal* parameter sets found in part two, we choose the ones yielding minimal *SSQ* = *SSQ*_*V*_ + *SSQ*_*S,E*_, as the overall optimal parameter set for the given patient.
4. To obtain confidence intervals for the optimal parameter estimates *p*_*opt*_ for each patient, we employ a bootstrapping technique. We assume that the best fit parameters yield the true dynamics, and that any discrepancy from the data is due to measurement errors. First, we calculate the residuals

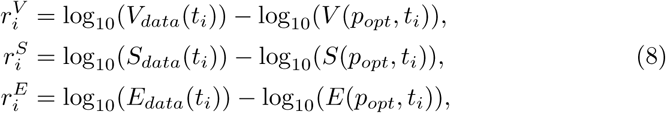

between simulated functions and measured data at times *t*_*i*_ (*i* = {1, *…*, 8}). Next, we create 1000 data sets for the HBV DNA, HBsAg, and HBeAg data at times *t*_−1_, *…, t*_8_, where data at times *t*_−1_ and *t*_0_ are as before and data at the remaining times are obtained by adding a randomly drawn residual (with repetition) to the true value at each time, *i.e.*

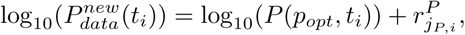

where *P* ∈ {*V, S, E*}, *i* = 1, *…*, 8, and *j*_*P,i*_ is drawn at random from {1, *…*, 8}. Lastly, for each data set, we find a new set of optimal parameters by using *Matlab’s fmincon* with initial parameter guess *p*_*opt*_ to minimize *SSQ*_*V*_ and *SSQ*_*SE*_, as described in (2.). This yields 1000 sets of parameters (one for each data sets), and the confidence intervals on the optimal parameters *p*_*opt*_ are obtained as the ranges from the 2.5th percentiles to the 97.5th percentiles of the 1000 parameter values.

## Results

### Parameter estimates

The best parameter estimates, the respective errors (*SSQ*) and the the 95% confidence intervals obtained by bootstrapping, are given in table 2. Numerical solutions for each population versus data are shown in Fig. 1 (see also Figs. 2, 3, and 4 for zoomed in results). Table 3 gives the parameters obtained from equilibrium conditions (6).

**Table 2.** Estimated parameters, fit errors, and confidence intervals.

| | $g$<br>($\times 10^{-2}$)<br>$d^{-1}$ | $c$<br>$d^{-1}$ | $d_S$<br>$d^{-1}$ | $d_E$<br>$d^{-1}$ | $1 - \epsilon$<br>($\log_{10}$) | $EC_2$<br>($\log_{10}$)<br>mg/kg | $EC_3$<br>($\log_{10}$)<br>mg/kg | $k_{eo}$<br>($\times 10^{-2}$)<br>$d^{-1}$ | SSQ |
| --- | --- | --- | --- | --- | --- | --- | --- | --- | --- |
| 703 | 4.6 | 1.33 | 0.12 | 1.52 | -3.8 | -3.39 | -3.49 | 4.69 | 1.03 |
| 703 | (3.7,5.8) | (1.3,1.6) | (0.11,0.16) | (1.3,1.7) | (-4,-3.6) | (-3.6,-3.2) | (-3.6,-3.4) | (4,5.4) |  |
| 704 | 0.3 | 9.27 | 0.6 | 1.35 | -2.94 | -3.25 | -2.96 | 9.81 | 1 |
| 704 | (0,1.4) | (6.8,11) | (0.5,0.7) | (1.1,1.5) | (-3,-2.7) | (-3.4,-3.2) | (-3,-2.8) | (8.5,11.2) |  |
| 708 | 2.54 | 1.24 | 0.25 | 0.6 | -3.67 | -3.39 | -2.61 | 6.43 | 1.42 |
| 708 | (0.6,5.1) | (1.1,2.5) | (0.2,0.3) | (0.5,0.8) | (-4.2,-3.3) | (-3.6,-3.3) | (-2.7,-2.5) | (5.2,7.7) |  |
| 710 | 4.48 | 1.87 | 0.15 | 0.37 | -3.43 | -3.73 | -3.02 | 8.3 | 1.48 |
| 710 | (2.9,6) | (1.3,2.4) | (0.1,0.18) | (0.2,0.5) | (-3.7,-3.1) | (-4.2,-3.4) | (-3.2,-2.8) | (5.9,11) |  |
| 711 | 2.41 | 3.12 | 0.21 | 1.4 | -4.13 | -3.14 | -2.8 | 5.55 | 0.78 |
| 711 | (1.4,3.2) | (2.8,3.3) | (0.2,0.24) | (1.1,1.7) | (-4.3,-3.9) | (-3.2,-3) | (-2.9,-2.7) | (4.7,6.4) |  |
| MEAN | 2.87 | 3.37 | 0.27 | 1.05 | -3.59 | -3.38 | -2.98 | 6.96 | 1.14 |
| MEDIAN | 2.54 | 1.87 | 0.21 | 1.35 | -3.67 | -3.39 | -2.96 | 6.43 | 1.03 |
| SD | 1.77 | 3.38 | 0.19 | 0.52 | 0.44 | 0.22 | 0.33 | 2.08 | 0.3 |

**Table 3.** Parameters obtained from fitted parameters in table 2, under equilibrium conditions defined by eq (6).

| | $p$ | $\alpha$ | $p_S$<br>($\times 10^{-3}$) | $p_E$<br>($\times 10^{-4}$) |
| --- | --- | --- | --- | --- |
| 703 | 1.99 | 1206.9 | 1.54 | 2.48 |
| 704 | 1.22 | 861.66 | 0.85 | 1.33 |
| 708 | 0.67 | 613.66 | 1.65 | 0.99 |
| 710 | 2.8 | 1569.85 | 1.63 | 1.77 |
| 711 | 9.37 | 4526.23 | 1.77 | 1.58 |
| MEAN | 3.21 | 1755.66 | 1.49 | 1.63 |
| MEDIAN | 1.99 | 1206.9 | 1.63 | 1.58 |
| SD | 3.54 | 1590.21 | 0.37 | 0.56 |

**Fig 1.**
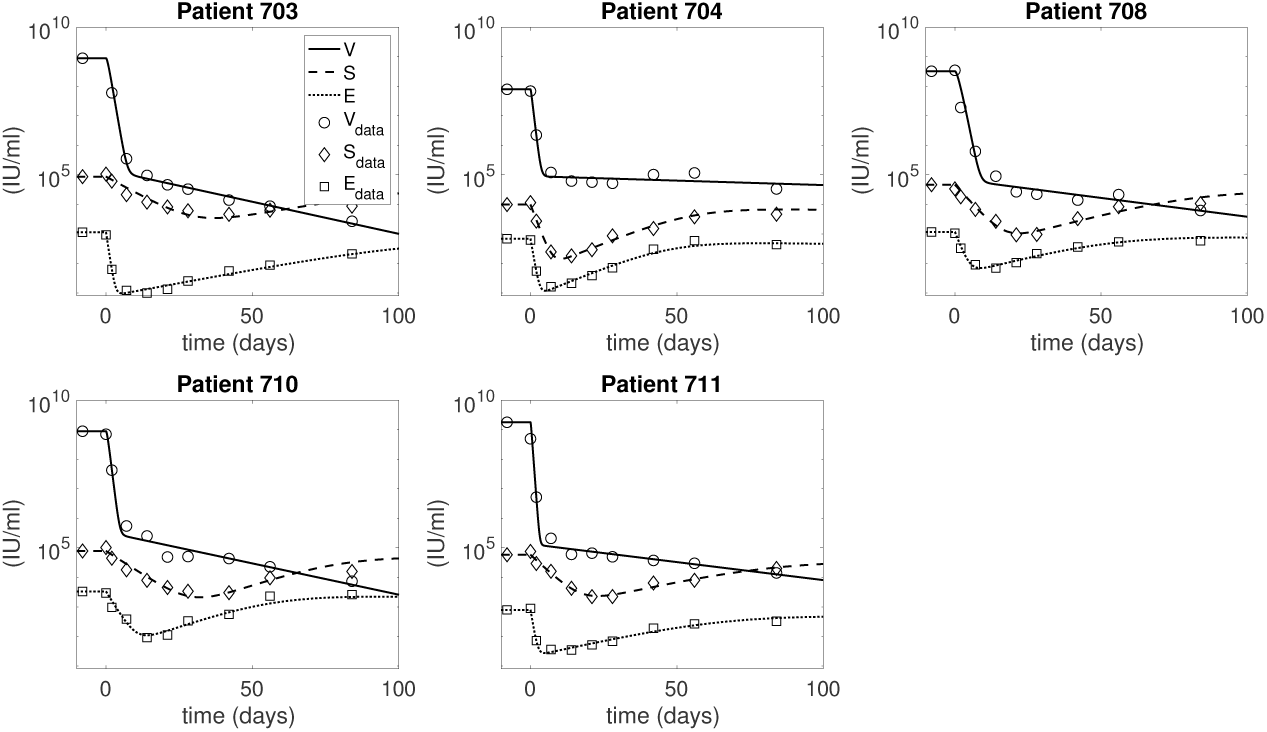
HBV DNA, HBsAg, and HBeAg dynamics over time as given by model (5) (solid curves) versus data (circles). The parameters are given in tables 1 and 2.

**Fig 2.**
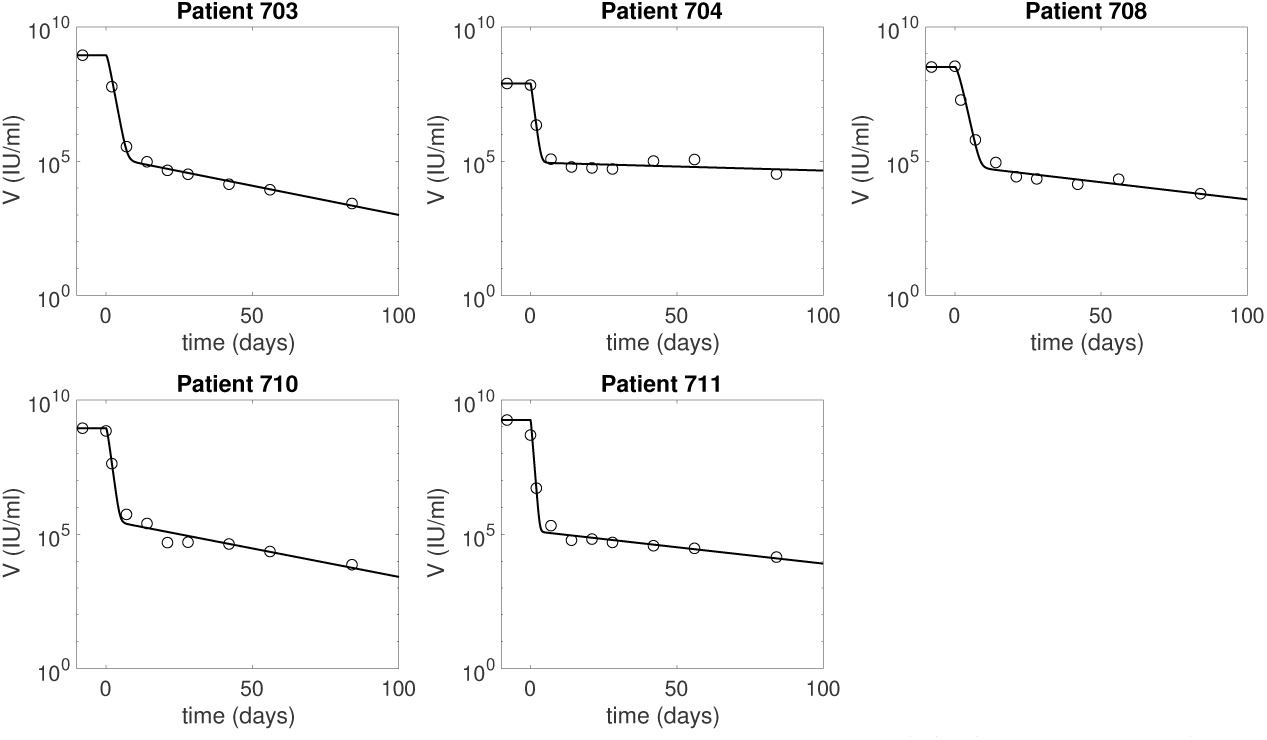
HBV DNA dynamics over time as given by model (5) (solid curves) versus data (diamonds). The parameters are given in tables 1 and 2.

**Fig 3.**
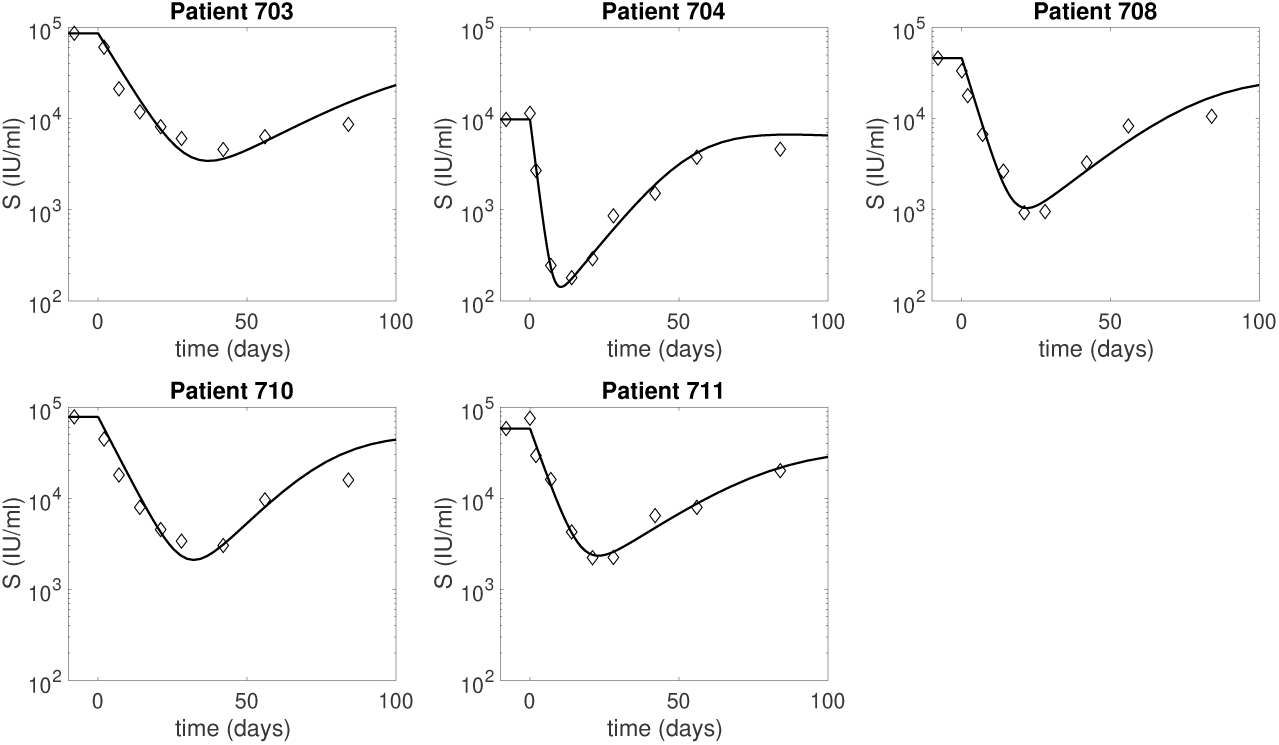
HBsAg dynamics over time as given by model (5) (solid curves) versus data (diamonds). The parameters are given in tables 1 and 2.

**Fig 4.**
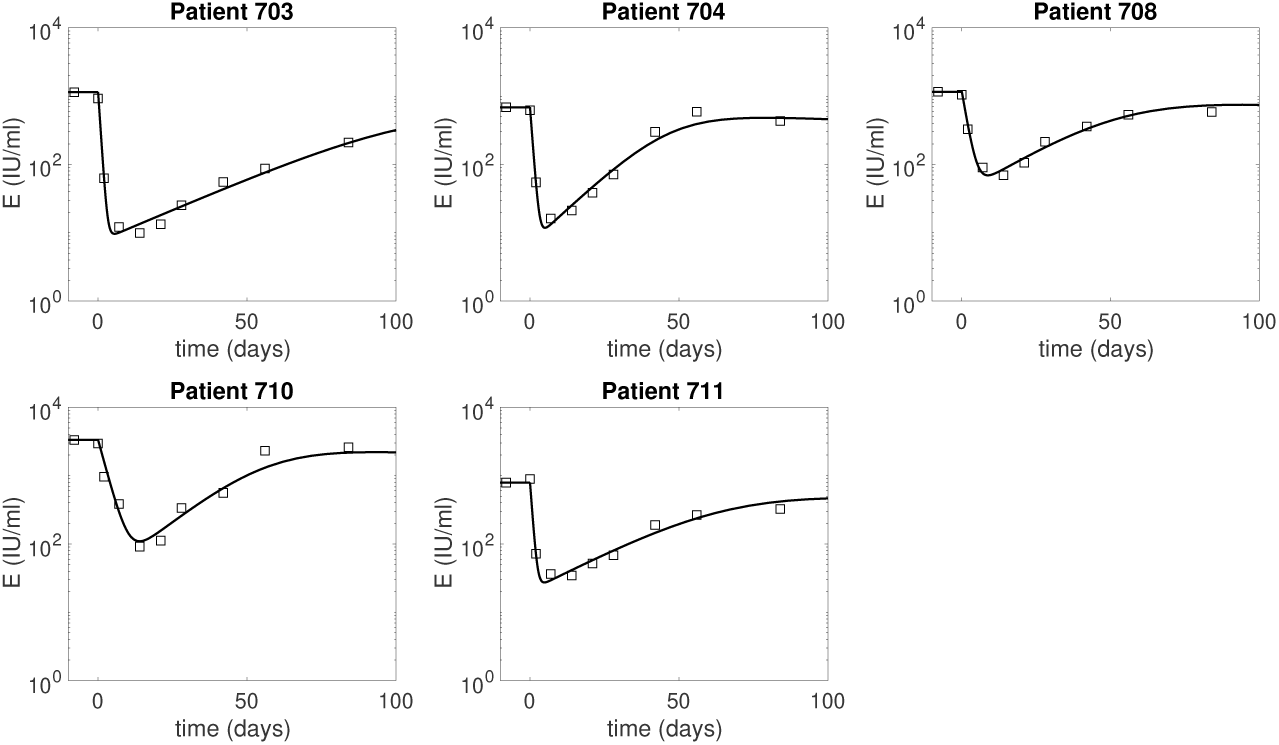
HBeAg dynamics over time as given by model (5) (solid curves) versus data (diamonds). The parameters are given in tables 1 and 2.

Previously reported virus clearance rates range from 0.69 per day [20, 28, 52] to 21.7 per day [53]. We estimate average virus clearance rates among the five patients *c* = 3.37 ± 3.38 per day, corresponding to average life-spans of 7.1 hours. The fastest free virus clearance rate, *c* = 9.27 per day (life-span of 2.6 hours), occurs in patient 704, who has the lowest pre-treatment virus titer. Assuming 50% of hepatocytes are HBV-infected, we estimate an average intracellular HBV DNA release rate *p* = 3.21 ± 3.54 per day. Patient 711, who has the highest pre-treatment virus titer, has *p* = 9.37 per day, 2.9 times higher than the average. Under these estimates, the pre-treatment serum virus production rates, *pD*_0_, range between 301.5 and 1260 copies/(infected cell×day) for patients 703–710, similar to the 200–1000 copies/(infected cell×day) reported for acute HBV infection [57]. Patient 711, however, has a pre-treatment serum virus production rate, *pD*_0_ = 4216.5 copies/(infected cell × day), four times larger than in [57]. Intracellular HBV DNA synthesis rates are *α* = 1755.66 ± 1590.21 copies/(cell × day). As with the serum release rate, patient 711 has 2.6-times higher intracellular HBV DNA synthesis than the average, *α* = 4526.23 copies/(ml × day).

The reported half-life of circulating HBsAg in chronically infected patients is 6.7 days (with a standard deviation of 5.5 days) [54], which corresponds to HBsAg decay rates 0.057 *< d*_0,*S*_ *<* 0.58 per day. We estimate average HBsAg decay rates *d*_*S*_ = 0.18 ± 0.06 per day, corresponding to HBsAg life-span of 5.6 days for patients 703 and 708-711, and *d*_*S*_ = 0.6 per day, corresponding to HBsAg life-span of 1.7 days, for patient 704. The average clearance rates of circulating HBeAg *d*_*E*_ = 1.05 ± 0.52 per day, correspond to HBeAg life-spans ranging between 15.8 hours and 2.7 days, about one order of magnitude lower than those reported by Loomba et al. for HBsAg [54]. The decreased HBeAg life-span predicted by our model may be correlated with the emergence of immune events and/or mutation in the core/precore regions [39] during ARC-520 treatment. Since we have no data on these events, we did not account for them in our model. Production rates of HBsAg and HBeAg are estimated to be *p*_*S*_ = (1.49 ± 0.37) × 10^−3^ IU/(cell × day) and *p*_*E*_ = (1.63 ± 0.56) × 10^−4^ IU/(cell × day), respectively.

We estimate high efficacy rates, *ϵ*_*T*_ *>* 99.88%, for the combined entecavir and ARC-520 effects in blocking HBV DNA synthesis. The additional time-dependent inhibitory effect on intracellular HBV DNA synthesis is on average *g* = 0.029 ± 0.018 per day, significantly lower than the estimate of 0.13 per day in HBV infected mice with humanized livers treated with lamivudine, but similar to the estimate of 0.059 per day in mice treated with pegylated interferon-*α*-2a [40].

The estimated *k*_*eo*_ = 0.07 ± 0.021 per day, predicts slow transport of ARC-520 from plasma to liver. The half-maximal quantities are small, with average log_10_(*EC*_2_) = − 3.38 ± 0.22 and log_10_(*EC*_3_) =− 2.98 ± 0.0.33 for the ARC-520 effects on HBsAg and HBeAg, respectively. This implies that the effects of ARC-520 are long-lived, as suggested by Schluep et al. [51] who found that RNA inhibitors persist and induce antiviral effects for longer than the drug’s life-span.

### Pharmacokinetic-pharmacodynamic model dynamics

The predicted HBV DNA populations as given by model (5) for the estimated parameters follow a biphasic decay with short and sharp first phase corresponding to the removal of HBV DNA followed by long and slow second phase decay due to time dependent treatment induced inhibition of intracellular HBV DNA synthesis and infected cell loss. HBsAg and HBeAg decay at steep rates during the first 24.67 ± 10.2 and 7.64 ± 3.95 days, respectively. After reaching minimum values, on average 1.57 ± 0.19 and 1.6 ± 0.33 orders of magnitude smaller than their initial levels, HBsAg and HBeAg rebound (see Fig 3 and 4). Once the effects of ARC-520 have completely waned, HBsAg and HBeAg decay at rate *δ*.

For the estimated parameters, ARC-520 effects *η*_2_ and *η*_3_ given by model (4) increase from 0 to their maximum values during the first (ln(*k*_*eo*_) − ln(*d*))*/*(*k*_*eo*_ − *d*) = 1.04 ± 0.07 days. The effect of ARC-520 on HBsAg is similar for all patients, with maximal effect at day 1 (ranging between *η*_2_ = 0.986 and *η*_2_ = 0.998), which wanes to *η*_2_ = 0.5 in 1.8 to 3.4 months (see Fig 5a). The maximal effect of ARC-520 on HBeAg at day 1 ranges between *η*_3_ = 0.96 (patient 708) and *η*_3_ = 0.993 (patient 703) and wanes to *η*_3_ = 0.5 within 1.5 to 3.5 months (see Fig 5b). For both HBsAg and HBeAg, the effect of ARC-520 lasts longest in patient 703.

**Fig 5.**
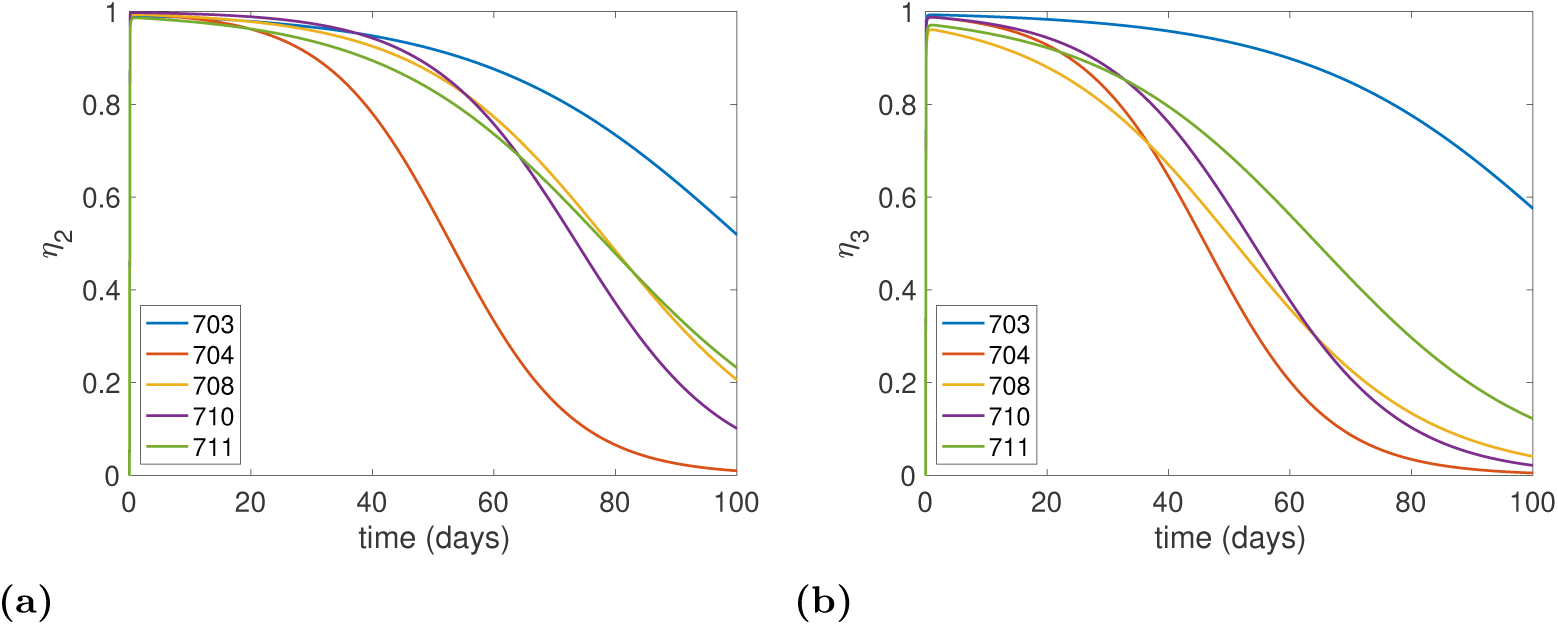
Efficacy of ARC-520 treatment over time as given by model (4) on (a) HBsAg production, and (b) HBeAg production. The parameters are given in tables 1 and 2.

### *In-silico* knockout experiments

We are interested in understanding the individual and combined effects of ETV and one-dose of ARC-520 on the dynamics of HBV DNA, HBsAg and HBeAg as given by model (5). We consider the following about the combined ETV and ARC-520 effects on reducing intracellular synthesis, *ϵ*_*T*_ : we either attribute it to ETV alone, 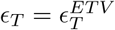; or split it between the two effects, 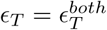. Using the parameters obtained from fitting the combination therapy model (5) to the Heparc-2001 clinical trial data [19], we conduct *in silico* experiments to determine how the dynamics change under: *in silico* monotherapy with entecavir, described by *η*_*i*_(*t*) = 0 for *i* = 2, 3, *g* ≠ 0, and 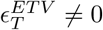; and combined entecavir and ARC-520 treatment, described by *η*_*i*_(*t*) ≠ 0 for *i* = 2, 3, *g* ≠ 0, and *ϵ*_*T*_ ≠ 0 (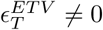, 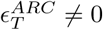, and 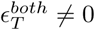) obtained through data fitting.

When we investigate *in silico* ETV monotherapy targeting HBV DNA intracellular synthesis, 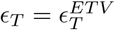, we can analytically derive the solutions of model (5) by considering *η*_2_ = *η*_3_ = 0. *g* ≠ 0, and 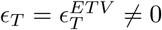. The infected cell population becomes *I*(*t*) = *I*_0_*e*^−*δt*^, the intracellular HBV DNA:

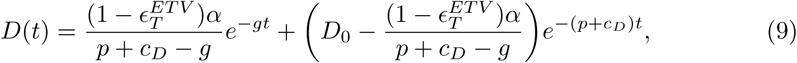

and extracellular HBV DNA:

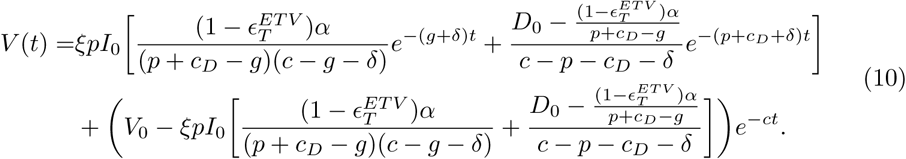

The equations for HBeAg is given by:

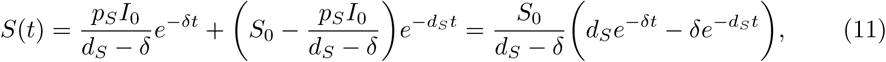

and for HBeAg is given by:

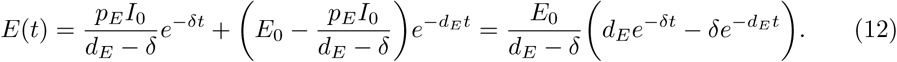

Note that both *S*(*t*) and *E*(*t*) are independent of *ϵ*_*T*_. HBV DNA follows a biphasic decay with short and sharp first phase corresponding to the removal of free virus followed by a slow second phase decay due to time dependent treatment induced inhibition of intracellular HBV DNA synthesis and removal of infected cells (see Fig 6, dashed curves). Serum antigen levels remain elevated for all three populations (see Fig 7 and 8, dashed curves).

**Fig 6.**
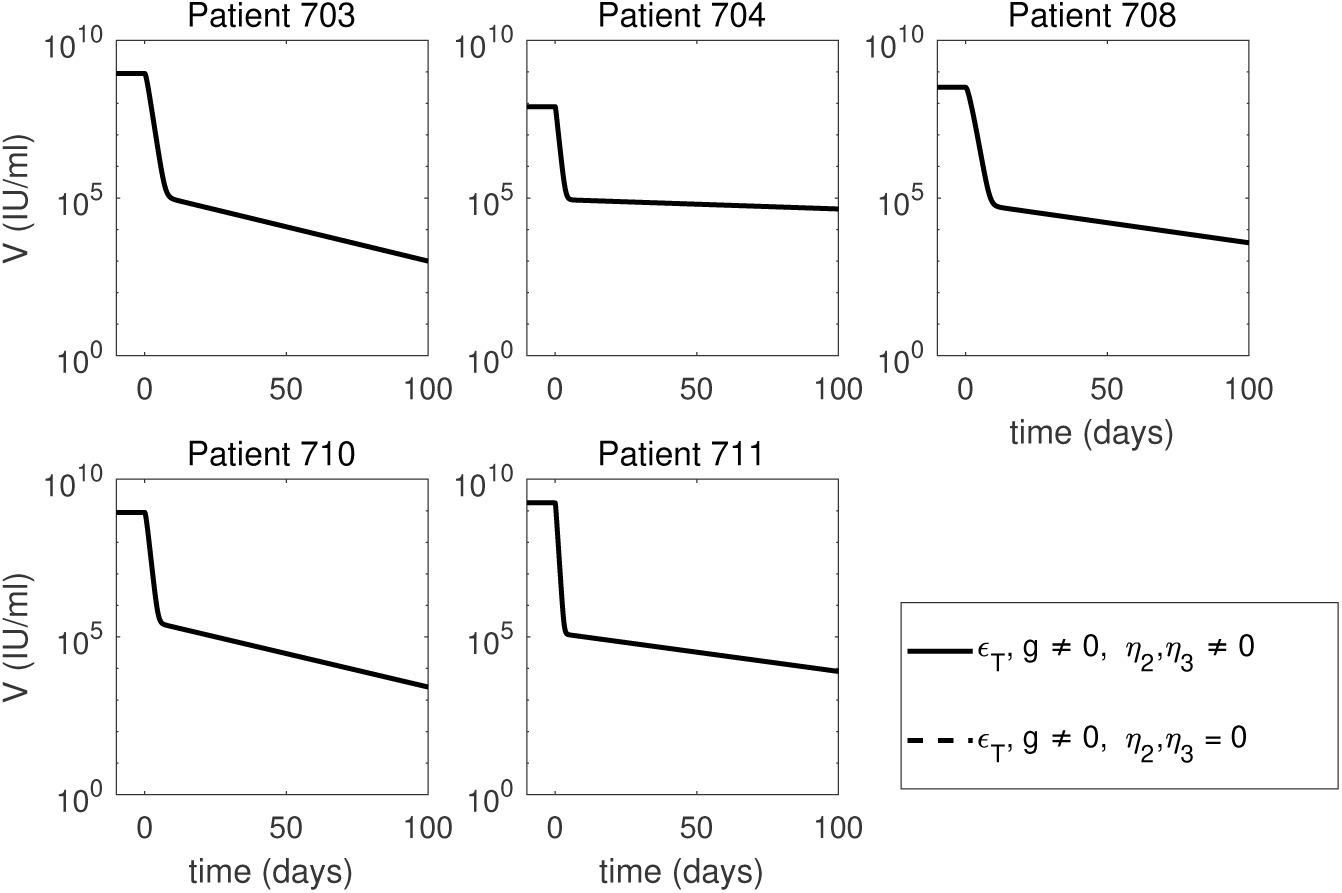
Short-term HBV DNA dynamics under ETV monotherapy (dashed curves), and combined ETV and ARC-520 therapy (solid curves), as given by model (5). Parameters are given in tables 1 and 2. Additionally, *g* ≠ 0, 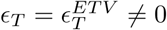 and *η*_2_(*t*) = *η*_3_(*t*) = 0 for ETV monotherapy. Note that both axes are plotted on log scale and that the two graphs overlap.

**Fig 7.**
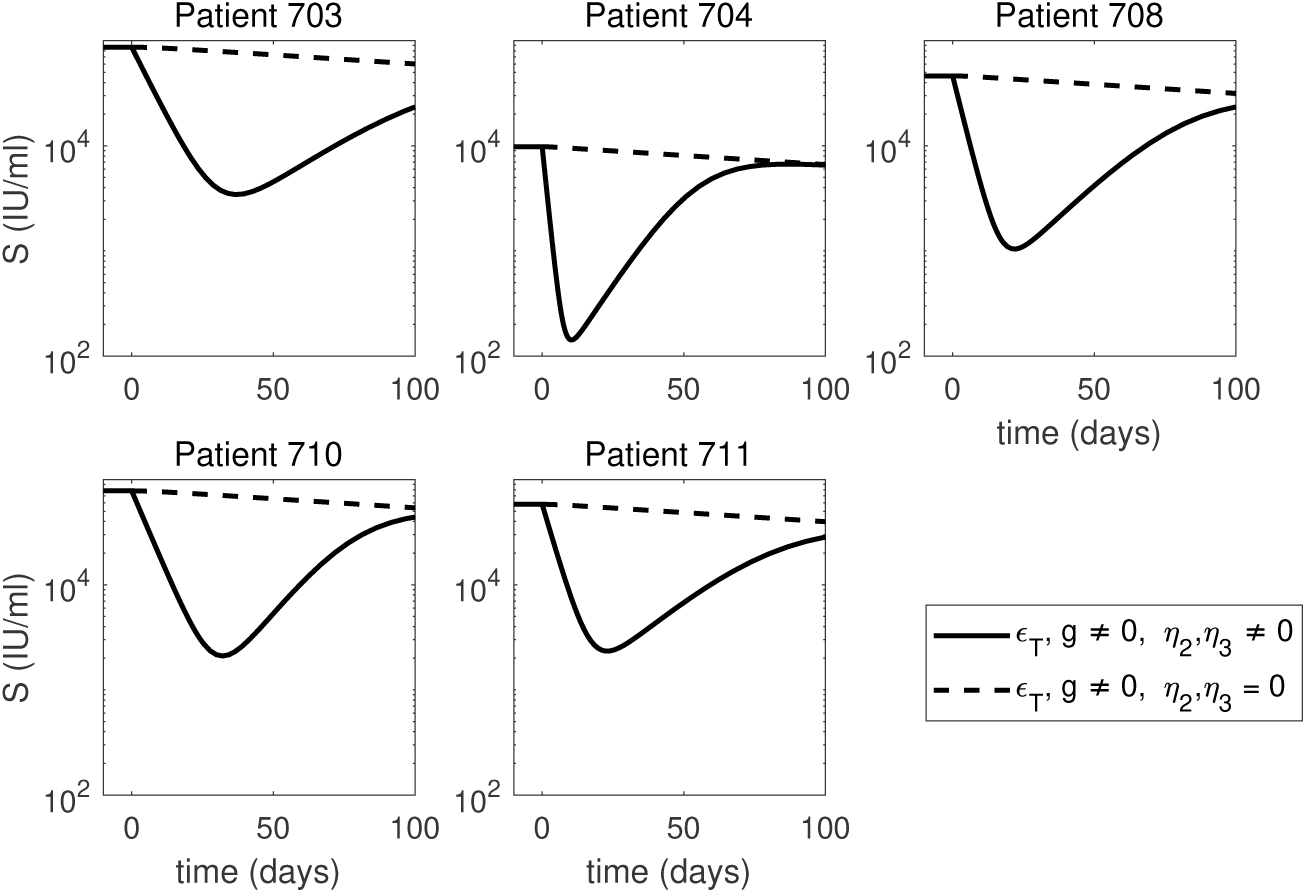
Short-term HBsAg dynamics under ETV monotherapy (dashed curves), and combined ETV and ARC-520 therapy (solid curves), as given by model (5). Parameters are given in tables 1 and 2. Additionally, *g* ≠ 0, 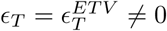 and *η*_2_(*t*) = *η*_3_(*t*) = 0 for ETV monotherapy. Note that both axes are plotted on log scale.

**Fig 8.**
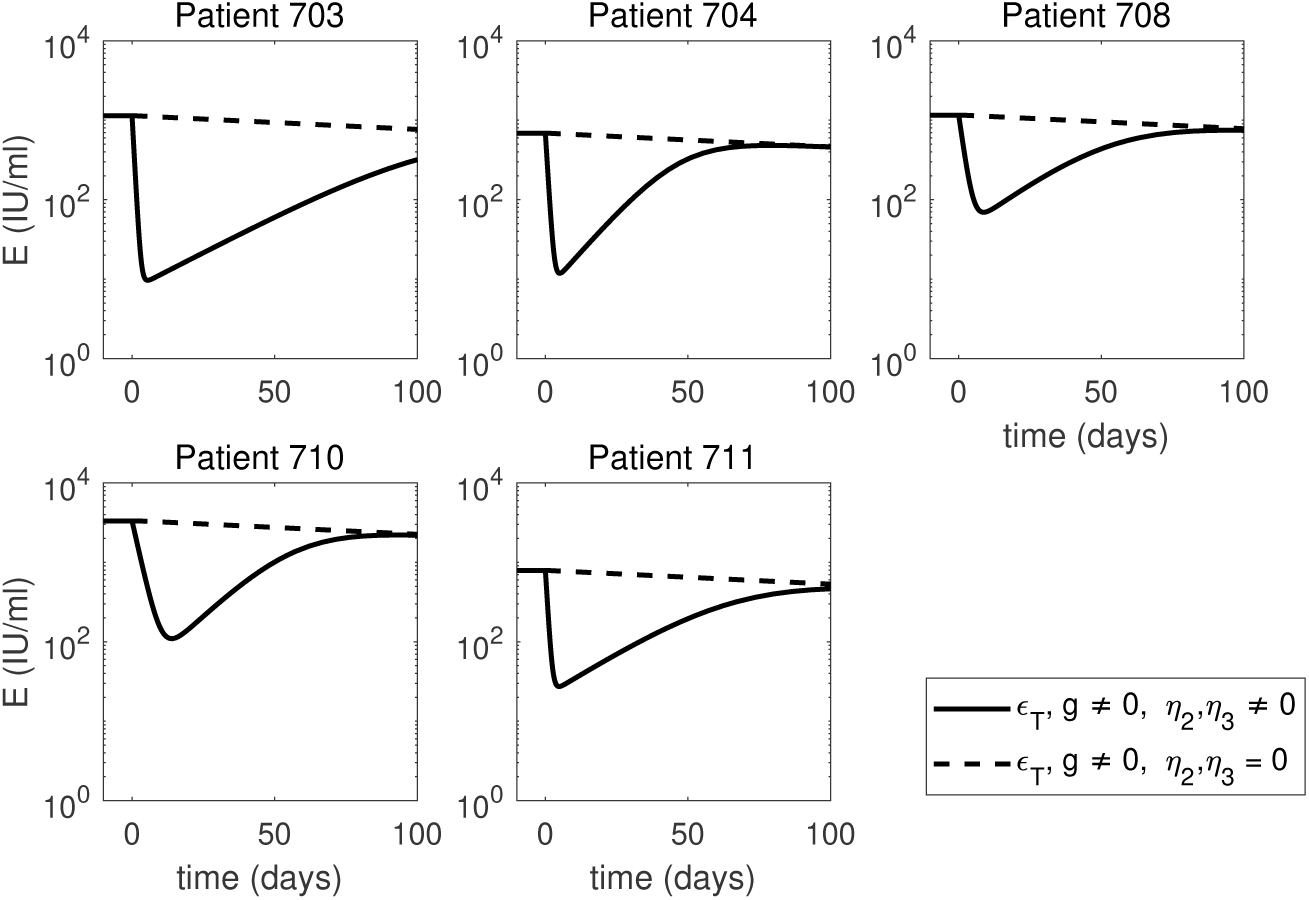
Short-term HBeAg dynamics under ETV monotherapy (dashed curves), and combined ETV and ARC-520 therapy (solid curves), as given by model (5). Parameters are given in tables 1 and 2. Additionally, *g* ≠ 0, 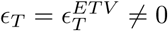 and *η*_2_(*t*) = *η*_3_(*t*) = 0 for ETV monotherapy. Note that both axes are plotted on log scale.

When we consider that the treatment that blocks intracellular HBV DNA synthesis, *ϵ*_*T*_, comes from both ETV and ARC-520, we recover the solutions of model (5) for combination therapy given by *η*_2_ = *η*_3_ ≠ 0, *g* ≠ 0, and 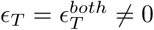. Both HBsAg and HBeAg decay at a steep rate during the first 22.7 ± 8.5 and 7.6 ± 4.1 days, respectively. After reaching minimum values, on average 1.5 ± 0.2 and 1.6 ± 0.4 orders of magnitude smaller than their initial levels, HBsAg and HBeAg rebound to their respective ETV monotherapy levels (see Fig 7 and 8, solid curves).

### Sensitivity of model predictions with respect to changes in the infected cell population’s initial condition

Previous estimates for the percentage of HBV-infected hepatocytes vary between 18 ± 12% in chronic HBsAg carriers [45, 46] and 99% in acute infections [21, 47]. We have derived our results by assuming that during chronic HBeAg-positive cases half of the liver is infected. Here, we investigate how changes in the size of the initial infected cell population alter our predictions. Analytical investigations show that the dynamics of the viral proteins HBsAg and HBeAg are not influenced by the initial size of the infected cell population, *I*_0_. After treatment initiation *I*(*t*) = *I*_0_*e*^−*δt*^, and *p*_*S*_ = *d*_*S*_*S*_0_*/I*_0_ and *p*_*E*_ = *d*_*E*_*E*_0_*/I*_0_ (based on the equilibrium assumption (6)). Therefore, the equations for *S* and *E*:

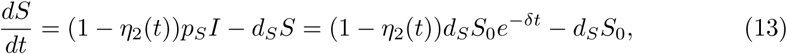

and

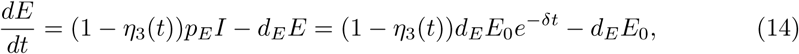

are independent of *I*_0_. Moreover, for *p* = *cV*_0_*/*(*ξD*_0_*I*_0_) and *D*_0_ = 225*/*(*I*_0_*/T*_*max*_) we find that intracellular HBV DNA *D* depends on *I*_0_ (see Fig 9) but HBV DNA in serum does not.

**Fig 9.**
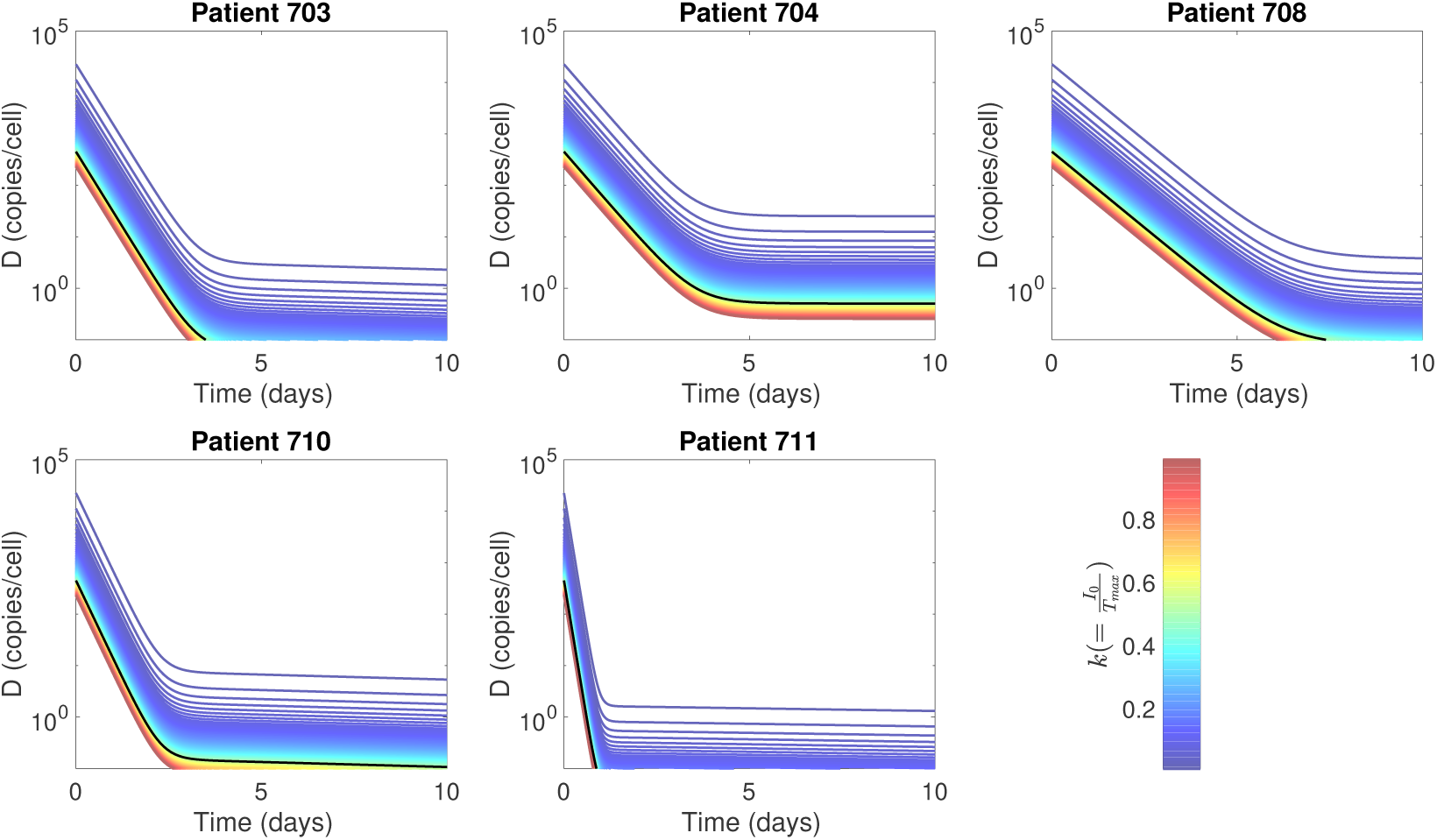
Intracellular HBV DNA dynamics of model (5) for 0.01 *< k <* 0.99 and *I*_0_ = *kT*_*max*_. Solid black lines show the dynamics for *I*_0_ = 0.5*T*_*max*_, which was used in data fitting. Other parameters used are given in tables 1, 2, and *D*_0_ = 225*/*(*I*_0_*/T*_*max*_).

### Long-term predictions and the need for uninfected hepatocyte dynamics

We assumed that infected hepatocytes have a life-span of 250 days. In this section, we are relaxing this assumption and investigate long-term HBV DNA and HBsAg dynamics when increased hepatocyte loss (due to either drug toxicity, or immune-mediated killing) is being considered. When we model it by increasing the infected cell death rate *δ* in (5) we obtain the following: long-term dynamics of *S* and *E* under ETV monotherapy predict that HBsAg decreases below 1 IU/ml 5.32 ± 0.54 months for *δ* = 7 × 10^−2^ per day, 4.21 ± 0.35 years for *δ* = 7 × 10^−3^ per year, and 7.35 ± 0.61 years for *δ* = 4 × 10^−3^ per day, following the initiation of therapy. Since ETV and other nucleoside analogues do not trigger cccDNA removal (and consequently HBsAg and HBeAg removal), the fast loss of HBsAg predicted by model (5) for higher killing rates *δ* is not realistic. In this section, we include the dynamics of uninfected and infected cell populations and investigate changes in predictions for increased killing rate *δ* We incorporate uninfected hepatocytes *T* which get infected by free virus at rate *β*, as modeled previously [21, 34, 58]. Note that we ignore the age of the infection and assume that once a cell becomes infected, it is producing virus (for a PDE model extension in a hepatitis C virus infection, see [59]). Both uninfected and infected hepatocytes proliferate according to a logistic term with maximal growth rate *r*_*T*_ and *r*_*I*_ and carrying capacity *T*_*max*_. In chronic HBV infections, cccDNA persist under long-term nucleoside analogues treatment [60]. Since the average cccDNA number of untreated HBeAg positive patients is 2.58 copies per infected cell [49], infected hepatocytes may have two infected offsprings. On the other hand, it has been suggested that cccDNA is destabilized by cell division or even lost during mitosis [60]. We account for this by assuming that a fraction Φ of proliferating infected hepatocytes have one infected and one uninfected offspring, and the remaining infected hepatocytes have two infected offsprings. The new model is given by:

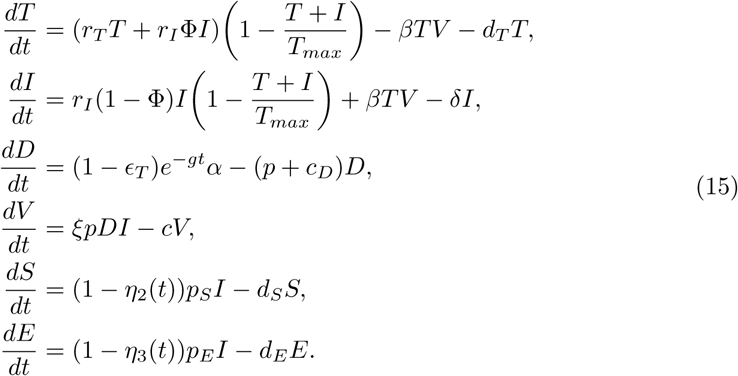

Liver regenerates rapidly after injury. To account for fast proliferation during chronic disease, we assume that hepatocytes’ maximum proliferation rate is *r*_*T*_ ≤ 1 per day, and *r*_*I*_ = 1 per day, corresponding to doubling time of (up to) 16 hours [21, 61]. The infectivity rate is at the lower end of previously fitted values [11], *β* = 10^−9^ IU/(ml × day); we include a death rate for the uninfected hepatocyte population, *d*_*T*_ = 4 × 10^−3^ per day [62], identical to that in model (5); and set the fraction of infected hepatocytes that have one uninfected and one infected offspring to Φ = 0.05. Initial conditions of uninfected and infected hepatocytes are set such that the model is in equilibrium prior to treatment with *D*_0_ = 450, and *V*_0_, *S*_0_, and *E*_0_ as in table 1. This leads to almost all hepatocytes being infected.

Without loss of generality, we investigate the dynamics for patient 703 under combination therapy for a continuum of *δ* values. Our hypothesis is that NA monotherapy cannot lead to HBsAg loss. In order to obtain infected cell persistence (under NA monotherapy), we need to decrease *r*_*T*_ (for a fixed *r*_*I*_ = 1) as *δ* increases (a *r*_*T*_ − *δ* threshold required for infected cells persistence is given in Fig 10). Therefore, HBsAg persistence under increased infected cell killing (as seen in NA treatment) may be explained by high ratio of infected to uninfected cell proliferation. Other events, such as HBV DNA integration, may also explain HBsAg persistence under infected cell (and potentially cccDNA) loss. This is especially true for HBeAg negative patients and NA experienced, HBeAg-positive patients.

**Fig 10.**
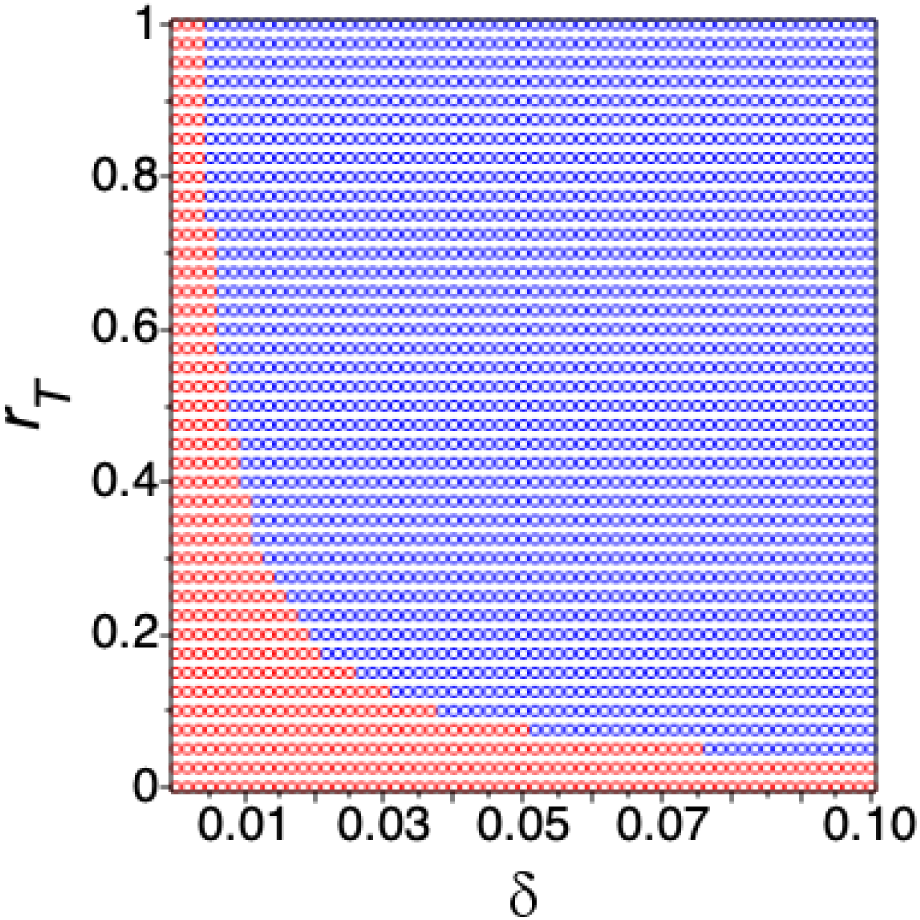
(*δ, r*_*T*_) ranges where infected cells given by model (15) are cleared (blue dots) or persist (red dots) under ETV monotherapy. Here *r*_1_ = 1 per day, *β* = 10^−9^ ml/(IU × day), *d*_*T*_ = 4 × 10^−3^ per day, initial conditions *T*_0_ and *I*_0_ are set such that the model is in chronic equilibrium in the absence of treatment. The other parameters are given in tables 1 and 2 for patient 703.

## Discussion

Reaching functional cure with current anti-HBV therapies in patients with chronic hepatitis B infection is hindered difficult by the lack of approved direct anti-HBsAg treatment and the presence of large numbers of HBsAg in the blood of infected patients [63, 64]. Therapies silencing viral translation through RNA interference technology [17, 19, 65], inhibiting HBsAg release via nucleic acid polymers [66–68], and inducing neutralization of HBsAg via specific antibodies [69, 70] have shown different levels of success [63, 64]. Understanding the relative effects in reducing HBV DNA, HBsAg and HBeAg titers of these new approaches alone, and in combination with traditional nucles(t)ide analogues, is particularly important in informing the development of new generation anti-HBsAg therapies.

To help in this endeavor, we developed mathematical models describing the HBV DNA, HBsAg and HBeAg in the presence of a silencing RNAi drug called ARC-520. We used the models and clinical trial data from treatment naive, HBeAg-positive patients that receive a one time ARC-520 injection and daily nucleoside analogue treatment with entecavir [19], to determine the efficacy of ARC-520 and nucleoside therapies on the short and long-term dynamics of HBV DNA, HBsAg, and HBeAg. To the best of our knowledge, we report for the first time that the time-dependent ARC-520 effects on HBsAg and HBeAg are more than 96% effective around day 1, and slowly wane to 50% in 1.8-3.4 months and 1.5-3.5 months, respectively. The combined ARC-520 and entecavir effect on HBV DNA is constant over time, with efficacy of more than 99.8%, which is similar to other nucleoside analogues trials.

A simplified version of the model, which ignored the dynamics of hepatocyte proliferation and infection, was sufficient to explain the short-term (about 100 days) dynamics observed in five patients in the current study. We found that one time injections with ARC-520 transiently reduce HBeAg and HBsAg titers, while daily nucleoside analogue treatments with entecavir reduce the viral load.

We modeled limited infected cell loss for the short-term dynamics. In the long-term, however, infected cells may die at faster rates, due to either drug toxic effects or increased immune killing. Lowering infected hepatocyte’s life-span to 100 (10) days, however, resulted in fast HBsAg removal, with decay below 1 IU/ml in 4.2 years (5.3 months). This loss, however, was in contradiction with clinical reports of low percentages of patients clearing HBsAg during long-term nucleoside analogues treatment [6], suggesting that more complex models are needed for long-term (several years) predictions. To determine under what conditions increased infected cells death does not spill over into unrealistic HBsAg and HBeAg loss under long-term nucleoside analogue therapy, we extended model (5) to include infected and uninfected cell dynamics. We assumed lower infected cells life-span (100 and 10 days), included division of both infected and uninfected populations, and determined that long-term HBsAg and HBeAg persistence under long-term HBV DNA clearance can be explained by high ratios of infected to uninfected division rates. Therefore, high ratio of infected to uninfected division rates, which correspond to the infection of the entire liver and may be indicative of scenarios where HBsAg seroclearance will not happen. Interestingly, us and others have associated high ratios of infected to uninfected division rates to triphasic HBV DNA decay under treatments with nucleoside analogues, a sign of suboptimal drug response [28, 30]. Whether infected hepatocytes indeed proliferate faster than uninfected hepatocytes remains under investigation.

While modeling results suggest that one-dose of ARC-520, in combination of daily entecavir, has limited long-term effects, we did not consider whether a transient reduction of HBsAg and HBeAg leads to the appearance of anti-HBs or anti-HBe antibodies, removal of immune-exhaustion, and eventual functional cure. Recent studies found that large levels of HBsAg might cause dysfunctional programming of HBsAg-specific B cells through persistent stimulation [71]. It has been suggested that therapeutic vaccines containing one (PreS2) or two (PreS1 or PreS2) envelope proteins together with serum HBsAg reducing drug therapies are needed in order to induce high levels of anti-HB antibodies, which may correlate with functional cure [72–74]. We ignored the level of immune modulation following RNAi based therapy, which is a model limitation, and therefore, we cannot say whether such effects were induced at higher rates during the transient HBsAg loss.

Our study has limitations. We only used the data on HBeAg-positive patients (cohort 7 in [19]) because they best responded to ARC-520 therapy and because they are the only studied cohort in which HBV DNA integration is not reported as a source of HBsAg production. Because of that, we excluded integration events from our mathematical model. A completely different modeling framework, that includes HBV DNA integration, is needed to investigate the ARC-520 effects on the HBsAg and HBeAg in the other cohorts containing patients who are either HBeAg-negative or NA-experienced and HBeAg-positive, with low cccDNA.

In conclusion, we developed a mathematical model and used it together with patient data, to estimate the time-dependent ARC-520 efficacies in blocking HBsAg and HBeAg productions. Additional data and theoretical efforts are needed to determine whether RNAi therapies have a feedback effect on the reversal of immune exhaustion, immunomodulatory immune responses, and potential functional cure.

## Supporting Information

Reference [40] was provided as a supplementary support.

## Acknowledgments

SK and SMC acknowledge funding from National Science Foundation grant No. 1813011. HD acknowledges funding from NIH grants no. R01AI144112 and R01AI146917. We thank the reviewers for their valuable comments.

## Author contributions statement

All authors conceived the study and performed the analyses. SK wrote the code. SK and SMC wrote the manuscript. All authors reviewed and revised the manuscript.

